# Approaches to maximize sgRNA-barcode coupling in Perturb-seq screens

**DOI:** 10.1101/298349

**Authors:** Britt Adamson, Thomas M. Norman, Marco Jost, Jonathan S. Weissman

## Abstract

Perturb-seq is a platform for single-cell gene expression profiling of pooled CRISPR screens. Like many functional genomics platforms, Perturb-seq relies on lentiviral transduction to introduce perturbation libraries to cells. On this platform, these are barcoded sgRNA libraries. A critical consideration for performing Perturb-seq experiments is uncoupling of barcodes from linked sgRNA expression cassettes, which can occur during lentiviral transduction of co-packaged libraries due to reverse transcriptase-mediated template switching. This problem is common to lentiviral libraries designed with linked variable regions. Here, we demonstrate that recombination between Perturb-seq vectors scrambles linked variable regions separated by 2 kb. This predicts information loss in Perturb-seq screens performed with co-packaged libraries. We also demonstrate ways to address this problem and discuss best practices for single-cell screens with transcriptional readouts.

## INTRODUCTION

Functional genomics efforts rely heavily on lentiviral transduction to introduce perturbation libraries into mammalian cells. Lentiviruses are powerful tools for manipulating host cells because they allow stable genomic integration of engineered DNA in both dividing and non-dividing cells, multiply or in single copy. However, experiments utilizing lentiviral systems must be designed carefully. This is because well-documented but underappreciated features of lentiviral biology produce inter- and intramolecular rearrangements that can disrupt transgene fidelity by scrambling linked variable regions or deleting sequences between repeats (Sack et al., 2016; ter Brake et al., 2008). Recent efforts to evaluate CRISPR perturbation libraries that carry such features have recently re-highlighted this constraint and, more generally, re-established the universal fact that mechanisms of reagent function are fundamental to experimental design (Feldman et al., 2018; Hill et al., 2018; Xie et al., 2018).

Most lentiviral transduction systems are based on HIV-1, an enveloped retrovirus that, upon entering a cell, converts its single-stranded RNA genome into double-stranded DNA that can integrate into host genomes (Hu and Hughes, 2012). Commonly, these transduction systems are composed of multiple expression vectors that separately encode viral elements, including so-called “transfer” plasmids that encode engineered viral genomes. These genomes carry the transgenes intended for host integration but also *cis-*acting viral elements necessary for virion packaging, RNA reverse transcription, and DNA integration (such as the viral long terminal repeats, LTRs) (Figure 1, 2). To minimize biosafety concerns, however, these genomes lack *trans*-acting HIV-1 genes. Instead, *trans*-acting elements, including HIV-1 reverse transcriptase (RT) and integrase (IN) as well as the G glycoprotein from vesicular stomatitis virus (VSV-G), are expressed from separate “packaging” and “envelope” plasmids that must be co-transfected into cells alongside transfer plasmids to produce infectious particles. Separation of viral elements in this manner ensures production of replication-incompetent virions. Importantly though, these engineered virions retain many biological features of wild type HIV-1. Notably, like HIV-1, they are pseudodiploid. This means that each infectious particle carries two single-stranded RNA molecules and, when produced in the presence of multiple distinct transfer plasmids, these can be heterotypic.

**Figure 1.**
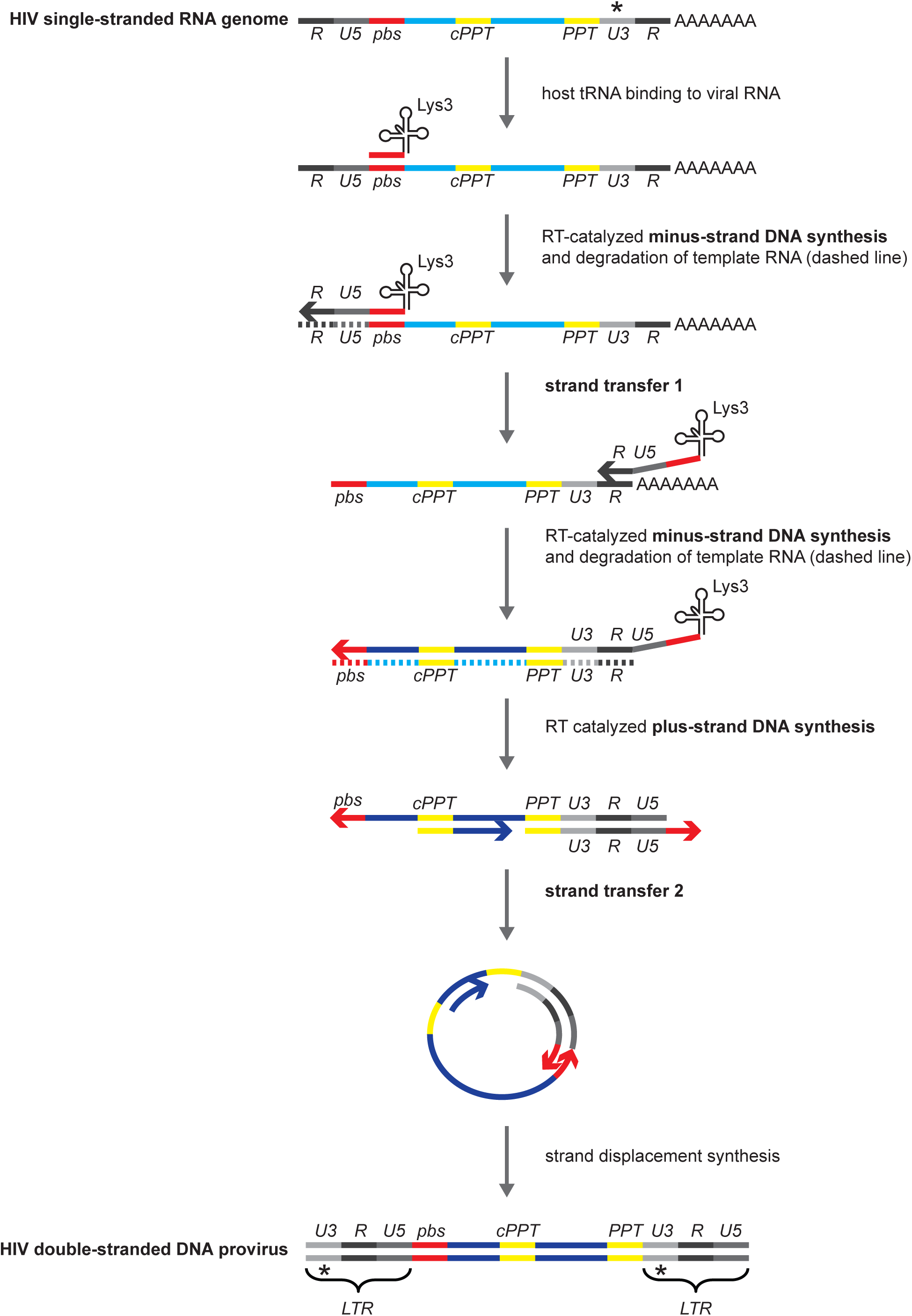
Reverse transcription of retroviral RNA into double-stranded DNA competent for host genome integration. Lys3, host tRNA. *pbs*, primer binding site. *PPT*, polypurine tract. c*PPT*, central polypurine tract. LTR, long terminal repeat. *, position of sgRNA expression cassette in CROPseq-Guide-Puro. RNA molecule is light blue.

**Figure 2.**
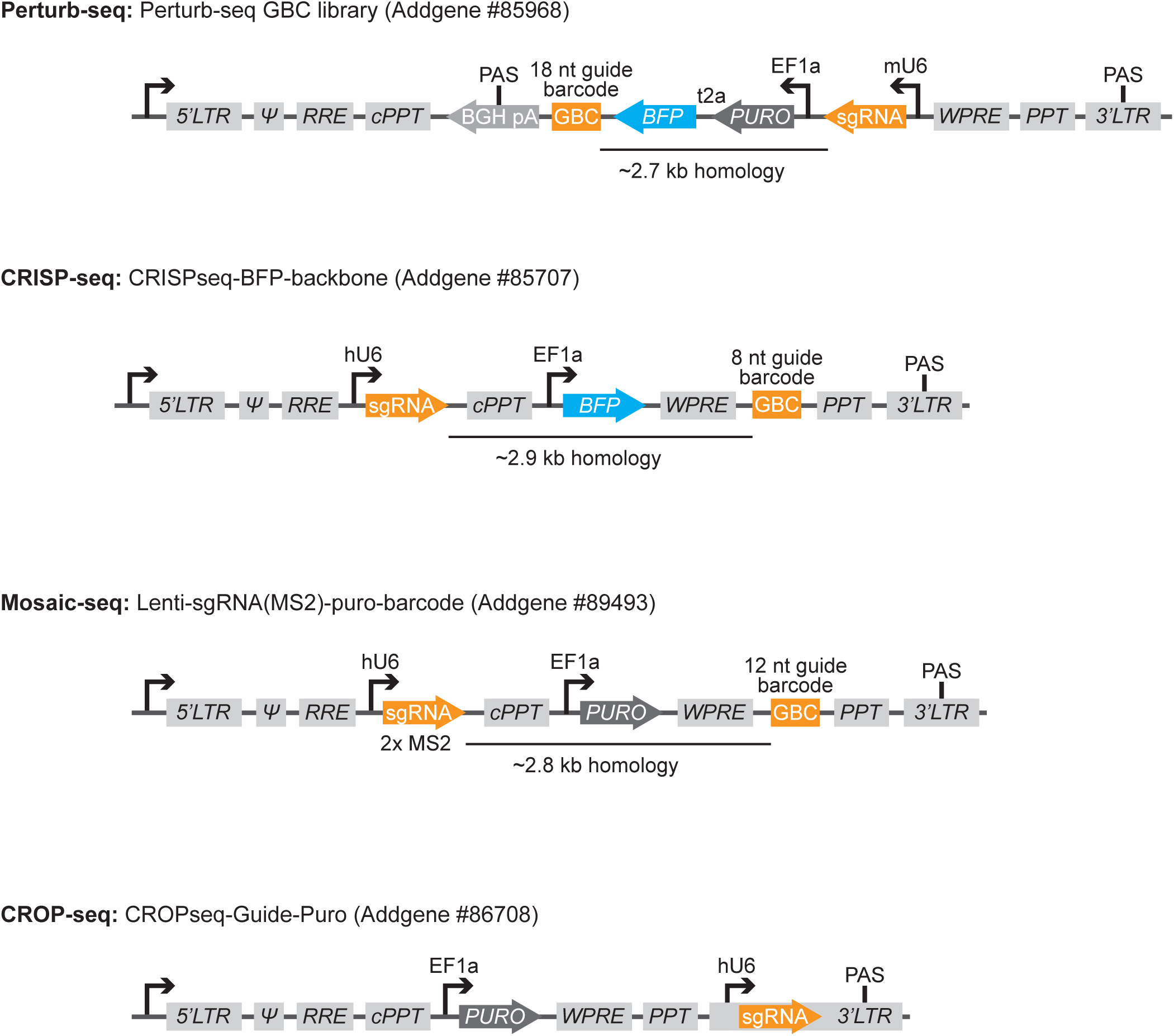
Published vector systems for scRNA-seq of CRISPR-based screens. Gray boxes indicate *cis*-acting elements: Ψ, Psi packaging element; RRE, Rev response element; WPRE, Woodchuck Hepatitis Virus Posttranscriptional Regulatory Element. BGH pA, Bovine Growth Hormone (BGH) polyadenylation signal and transcription termination sequence. BFP, blue fluorescent protein. PURO, puromycin resistance gene. GBC, guide barcode. PAS, polyadenylation signal sequence.

Given that engineered lentiviruses rely on retroviral biology for transgene delivery, one must consider wild type mechanisms of the viral life cycle, most notably HIV-1 reverse transcription (Figure 1), to understand how intra- and intermolecular rearrangements that can confound transduction arise. Reverse transcription of HIV-1 begins when a 5ʹ region of the RNA genome, called the primer binding site (*pbs*), anneals to a host tRNA molecule. This molecule acts as a primer to initiate a tract of RT-catalyzed DNA synthesis that copies the R-U5 region of the 5ʹ LTR. Facilitated by genome-flanking R repeats, this fragment then completes an obligatory intra- or intermolecular strand transfer (strand transfer 1) to the 3ʹ end of genomic RNA. This strand transfer enables the resumption of minus-strand DNA synthesis and notably completion of one U3-R-U5 LTR. As DNA synthesis proceeds, RNase H activity of the RT enzyme degrades template RNA. However, two purine-rich sequences, the polypurine tract (*PPT*) and the central polypurine tract (*cPPT*), remain intact to act as primers that initiate plus-strand DNA synthesis. Extension of the plus strand proceeds to incorporate 18 nucleotides templated by the host tRNA. These enable intramolecular annealing of the plus and minus strands (strand transfer 2) and subsequent extension that completes the double-stranded DNA provirus. Strand transfer 2 is strictly intramolecular and therefore linked LTRs (U3-R-U5) are identical (Yu et al., 1998). However, template switching occurs frequently during minus-strand DNA synthesis and thus can generate chimeric proviruses from heterotypic genomes. Indeed, markers separated by 1 kb in HIV-1 have been observed to segregate near-randomly (Rhodes et al., 2003).

Recently, we leveraged lentiviral transduction to deliver genetic perturbation libraries to mammalian cells in a manner that enabled pairing of CRISPR-based genetic screens with droplet-based single-cell RNA-sequencing (scRNA-seq) (Adamson et al., 2016; Dixit et al., 2016). This technique, called Perturb-seq, now allows multiplexed evaluation of perturbation phenotypes with single-cell resolution. Our first iteration of Perturb-seq relied on a straightforward library design that enabled single-cell identification of perturbations by coupling expression of CRISPR guide RNAs (sgRNAs) to unique guide barcodes (GBCs) (Figure 2). These variable elements were encoded approximately 2.7 kb apart within an engineered lentiviral genome, and for this reason, sgRNA-GBC uncoupling during transduction of co-packaged virions became a near certainty. We addressed and circumvented this limitation from the outset (Adamson et al., 2016; Dixit et al., 2016). However, work with similar systems, has recently re-emphasized template switching as a potential pitfall of Perturb-seq screens (Feldman et al., 2018; Hill et al., 2018; Xie et al., 2018). Moreover, perturbation calling by proxy from dual expression vectors, as in Perturb-seq, has become common among methodologies that pair scRNA-seq with functional genomics (Figure 2). These include CRISP-seq, Mosaic-seq, and CROP-seq (Datlinger et al., 2017; Jaitin et al., 2016; Xie et al., 2017). Here, we provide functional characterization of recombination during transduction of co-packaged Perturb-seq vectors and discuss methods to avoid this limitation. We also review current approaches for single-cell CRISPR screens and discuss best practices for these experiments.

## RESULTS

To investigate sgRNA-GBC uncoupling during transduction of co-packaged Perturb-seq vectors, we cloned two constructs from our standard backbone that enabled detection of recombinants (Figure 3A, B). These retain the basic design of Perturb-seq expression constructs; however, they encode linked markers that can be functionally distinguished by flow cytometry when transduced into GFP expressing cells. Specifically, BFP-sgGFP (pMJ002) encodes a blue florescent protein (BFP) approximately 2 kb away from an sgRNA targeting GFP (sgGFP), while mCh-sgNT (pBA613) encodes, in the same relative positions, mCherry florescent protein and a non-targeting sgRNA (sgNT) that does not deplete GFP. Using viruses produced in individual packaging reactions and therefore carrying homotypic genomes, we first compared co-transduction to individual transduction and observed no recombination of the linked markers (Figure 3C-E). However, when co-packaged viruses were transduced into cells (at infections rates that favored single copy integration) recombinants were produced at a high frequency (Figure 3C-E). Specifically, we observed that 21.8% of BFP+ cells were GFP+ compared to 5.4% of controls transduced with individually packaged viruses, indicative of 16.4% co-delivery with the unlinked sgNT, and 16.8% of mCherry+ cells were GFP-compared to 2% of controls transduced with individually packaged viruses, indicative of 14.8% co-delivery with sgGFP. Assuming equal availability of RNA genomes during viral packaging and random selection, which is consistent with HIV-1 (Lu et al., 2011), these data suggest ~30% marker exchange (Figure 3B). Notably, this result deviates from our initial expectation that, due to the ~2 kb of separating homology, markers would segregate randomly; however, our rate is consistent with a recent report that evaluated a similar system (Feldman et al., 2018).

**Figure 3.**
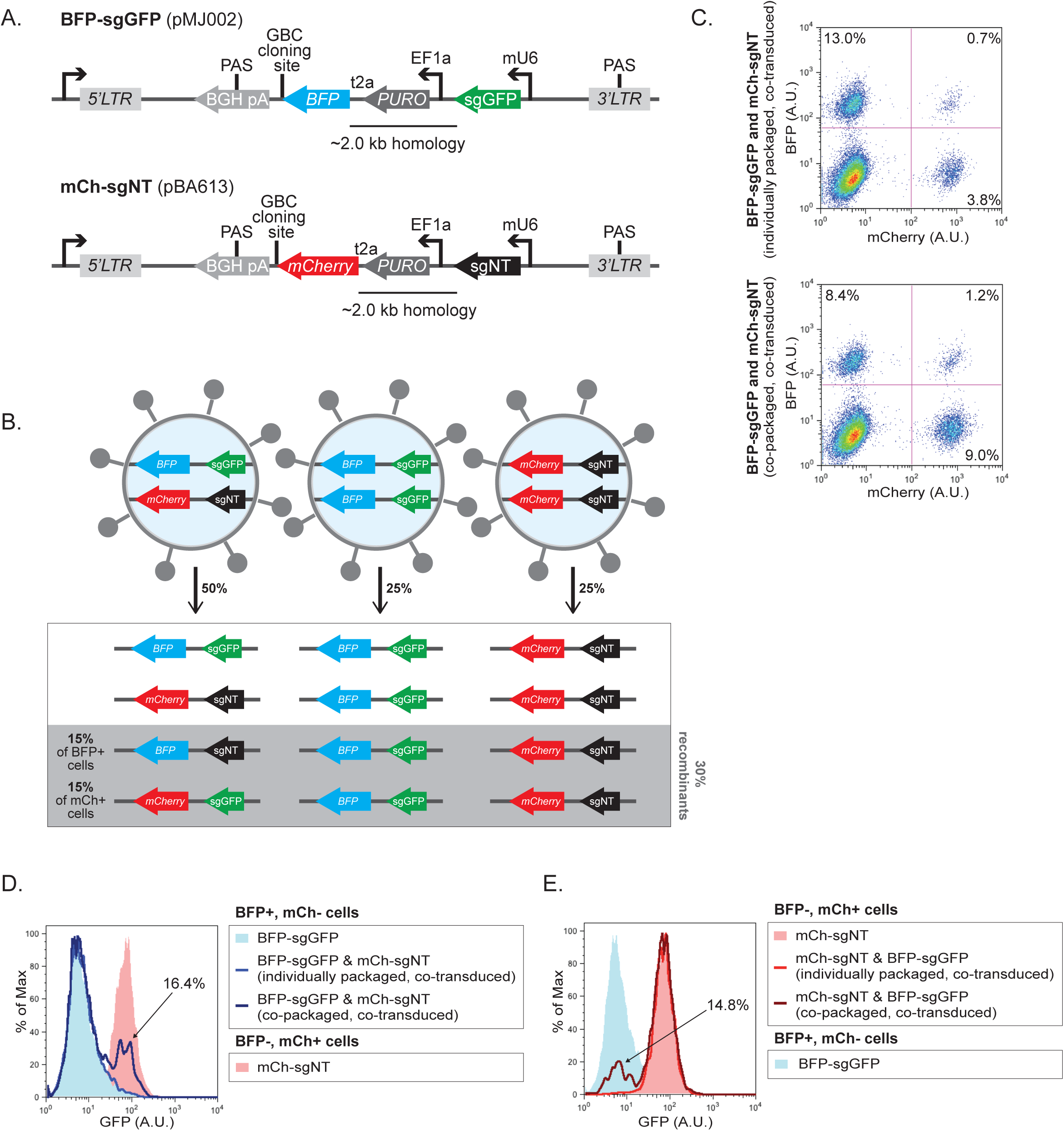
A. Schematic of BFP-sgGFP and mCh-sgNT constructs built to investigate recombination during transduction of co-packaged Perturb-seq vectors. **B.** Schematic of pseudodiploid viral genomes in virions (top) and resulting integrations in host DNA (bottom) possible after transducing cells with co-packaged BFP-sgGFP and mCh-NT. Products of recombination in gray box. Percentages included are those predicted for random genome packaging and 30% marker swapping. **C.** BFP and mCherry signals from cells transduced with BFP-sgGFP and mCh-sgNT using individually (top) or co-packaged (bottom) virions. Data from these samples also depicted in D and E. **D.** GFP expression in K562 cells carrying dCas9-BFP-KRAB (low background BFP signal) after transduction with the indicated constructs using viruses packaged as indicated. Data are presented for cells after gating for BFP+ and mCherry-(blue) or BFP- and mCherry+ (pink) as demonstrated in C. Percentage of observed recombinants from co-packaged condition indicated. **E.** GFP expression in K562 cells carrying dCas9-BFP-KRAB (low background BFP signal) after transduction with the indicated constructs using viruses packaged as indicated. Data and percentage of recombinants are presented as described in D.

Positioning the fluorescence marker genes in BFP-sgGFP and mCh-sgNT immediately upstream of the Perturb-seq GBC cloning site allowed us to quantify recombination events across a homology distance of ~2 kb. In Perturb-seq libraries, however, GBCs and sgRNAs are separated by a greater stretch of homology, ~2.7 kb (Figure 2). Previous work demonstrated that recombination frequencies increase with longer homologies between markers (measured up to 720 bp) (Sack et al., 2016). Therefore, our observations generate a lower bound of the sgRNA-GBC uncoupling frequencies that would be expected during transduction of co-packaged Perturb-seq libraries.

Promisingly, our results also suggest one straightforward strategy to ensure maintenance of sgRNA-GBC pairing in pooled Perturb-seq experiments: arrayed preparation of lentiviruses. We previously used this approach to conduct a pooled Perturb-seq experiment with 91 sgRNAs (Adamson et al., 2016). Although the arrayed approach is practical for experiments of moderate scale, alternative approaches that enable co-packaging while suppressing sgRNA-GBC uncoupling will also be greatly advantageous. One such approach is library co-packaging with so-called “decoy” genomes. These are genomes that lack homology to the sequence between linked variable regions and, when expressed at appropriate ratios, are expected to suppresses packaging of two Perturb-seq genomes into a single virion. This approach was recently piloted for a system similar to Perturb-seq (Feldman et al., 2018). Independently, we evaluated this approach for Perturb-seq. For this, we built a third transfer plasmid. This vector, lentiDecoy (pBA889), encodes neomycin under a doxycycline-inducible promoter and lacks homology with the sequence between markers in BFP-sgGFP and mCh-sgNT (Figure 4A). Promisingly, BFP-sgGFP and mCh-sgNT co-packaging with lentiDecoy (at a DNA balance of 4:4:92) achieved suppression of recombination between markers in the Perturb-seq vectors by ~4-fold, as determined by flow cytometry (Figure 4B,C). Moving forward, transfer plasmids rendered integration defective by mutations of the integrase recognition sequences (*att*) in the viral LTRs could be optimized for use as more inert decoys (Figure 4A). Indeed, recent work has shown that such a non-integrating decoy can be effective (Feldman et al., 2018).

**Figure 4.**
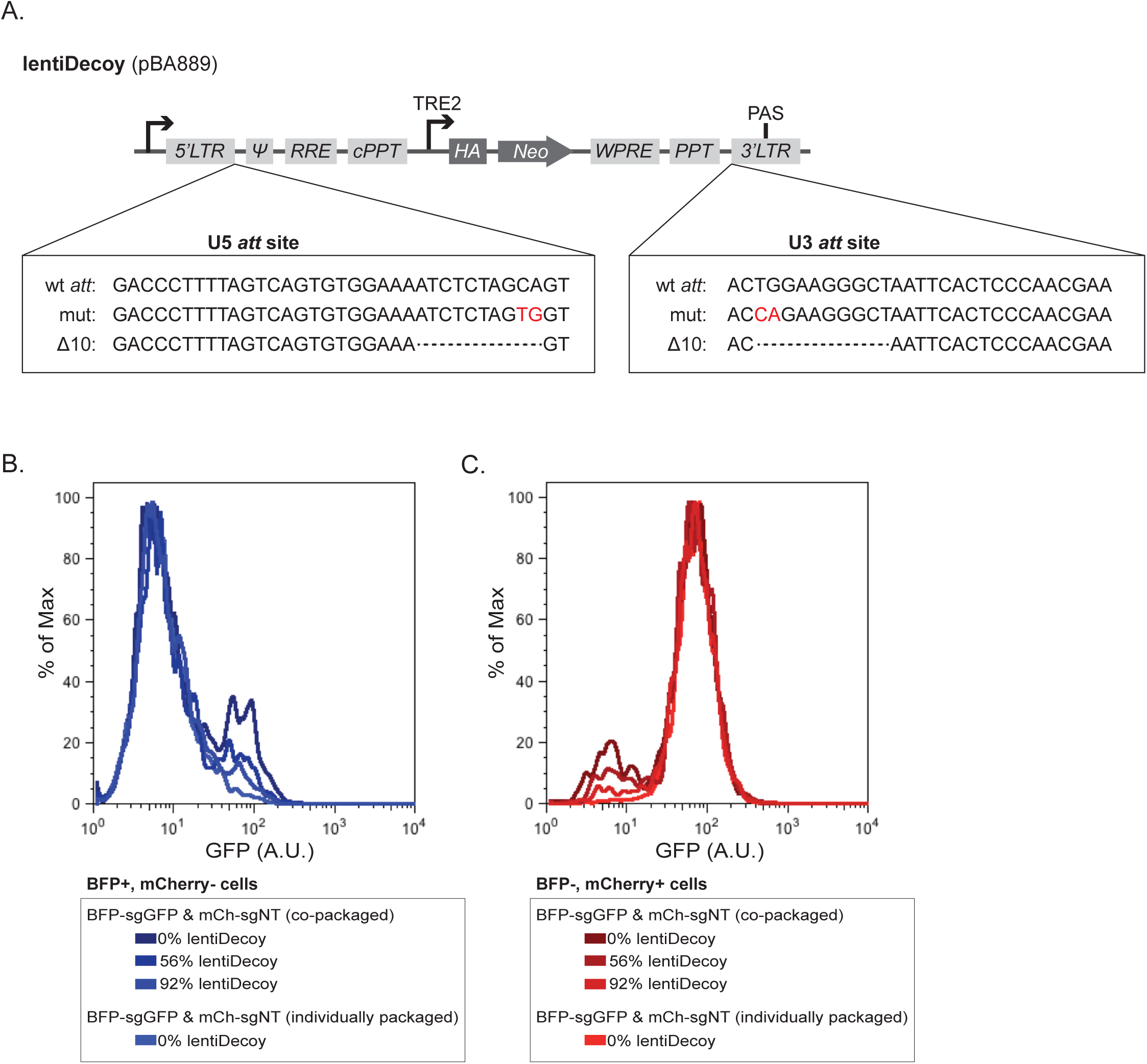
A. Schematic of lentiDecoy vector. Boxes include *att* site mutational strategies for rendering lentiDecoy integration deficient (Masuda et al., 1995). **B.** GFP expression in K562 cells carrying dCas9-BFP-KRAB (low background BFP signal) after transduction with the indicated constructs using viruses packaged as indicated. Data are presented for cells after gating for BFP+ and mCherry- (blue) or BFP- and mCherry+ (red). Traces for conditions without lentiDecoy are also depicted in Figure 3D. **C.** As in B. Traces for conditions without lentiDecoy are also depicted in Figure 3E.

## DISCUSSION

Single-cell RNA-sequencing (scRNA-seq) platforms typically capture the 3ʹ ends of polyadenylated transcripts and, when optimally expressed from RNA polymerase III (Pol III) promoters, sgRNAs are not polyadenylated. Perturb-seq screens thus rely on co-expression of polyadenylated barcodes (GBCs) for perturbation assignment. Here, we show that Perturb-seq expression constructs recombine during lentiviral transduction, which suggests that sgRNA-GBC uncoupling is likely to be pervasive in screens performed with co-packaged libraries. This concern is consistent with what is known about template switching in the lentiviral life cycle (Hu and Hughes, 2012) and with previous work that showed how uncoupling of linked variable regions can confound conventional functional genomics experiments performed with lentiviral libraries (Sack et al., 2016). More recently, *in silico* modeling of barcode uncoupling with Perturb-seq data and experimental evidence using a similar library design have demonstrated how this effect can diminish screen sensitivity (Hill et al., 2018). However, such uncoupling is unlikely to produce major spurious effects when left uncontrolled in highly complex Perturb-seq screens because uncoupled GBCs will mark diverse perturbations and thus any single misassigned behavior will be diluted within a guide assignment group. Nevertheless, to improve results, we suggest straightforward protocols that can circumvent or suppress uncoupling. These include arrayed library preparation and the use of “decoy” genomes.

We have previously used arrayed library preparation to prevent barcode uncoupling in pooled Perturb-seq screens (Adamson et al., 2016). For one of these experiments, we individually packaged 91 viruses from arrayed Perturb-seq vectors, titered each virus by test infection into target cells, and then mixed viruses with optimized representation for pooled transduction. Although relatively laborious, this procedure is readily applicable at the scale of hundreds of sgRNAs, and notably, most Perturb-seq experiments performed to date have been conducted at this scale, evaluating tens to hundreds of perturbations.

The choice to perform Perturb-seq experiments at modest scale reflects a practical reality: scRNA-seq can be prohibitively expensive. This limits tolerance for single-cell experiments that capture and sequence large numbers of uninformative or uninteresting cells, such as those with improper perturbation assignments. Importantly, this consideration has also kept the number of cells per perturbation low in Perturb-seq experiments. We previously found that 100-200 cells per perturbation is a good minimum for retaining transcript-level resolution of transcriptional effects and 10s of cells per perturbation retains expression signatures (Dixit et al., 2016). However, at these low cell numbers, maintaining even representation across complex perturbation libraries becomes a challenge, and this experimental hurdle is compounded for any perturbation that induces substantial growth defects.

Considering these challenges, arrayed preparation of Perturb-seq viruses, followed by individual titering in test infections and subsequent generation of balanced transduction pools, has multiple advantages. Principally, this approach is refractory to sgRNA-GBC uncoupling because it only generates virions with homotypic genomes. A second benefit comes from the fact that test infections enable the simultaneous quantification of viral titer and sgRNA effects on cell growth. This enables tight control over experimental coverage and buffering against dropout effects. In our hands, arrayed preparation followed by pooled transduction lead to a coefficient of variation of ~25% relative to intended representation at the end of the experiment, even with perturbations that are profoundly toxic (Adamson et al., 2016).

Of course, balanced reagent pooling can be performed at multiple points during complex library preparation, and use of decoy genomes to enable the use of co-packaged libraries without recombination concerns could enable earlier mixing. One strategy would be to pool individually cloned plasmids (based on molarity) for pooled viral preparation. Another would be to pool sgRNA-encoding oligos for batch cloning of expression vectors. However, when generated by array-based synthesis, variance in oligo representation can span an order of magnitude, and balancing procedures for either of these strategies may still be laborious. Buffering variation from cloning, for example, could require rounds of library construction, evaluation by sequencing, and recloning. Moreover, buffering perturbation dropout will always require *a priori* measurement of sgRNA effects on target cell growth. Therefore, although we (and others) are interested in these approaches for larger-scale experiments, the reduction in experimental effort they afford for moderate-scale experiments may be negligible.

Importantly, drawbacks of using decoy genomes in Perturb-seq experiments must also be considered. One of these is a drop in relative viral titer. If, for example, decoy genomes are expressed in packaging cells at a ratio of 9:1 with Perturb-seq genomes, then the vast majority (~81%) of virions produced from these cells will not carry a Perturb-seq genome, and assuming heterotypic genomes each have an equal chance of integrating, only 10% of produced viruses will deliver a perturbation. This is a 10-fold drop in effective titer. Of course, to deliver perturbations at representation, cells could be infected with more virions (higher overall multiplicity of infection, MOI). However, this presents concerns of spurious cell behaviors from superinfection and/or the genome integration of multiple proviruses. Notably, with an MOI of 0.1, only ~5% of infected cells are expected to receive more than 1 virion, but with an MOI of 1, multiple infections are expected in ~42% of infected cells. Alternatively, decoy prepared libraries could be transduced into more cells to maintain representation without high MOI. However, implementation of this approach may suffer from the practical difficulty of selecting a small minority of cells out of a large population for scRNA-seq. For example, only 5% of cells infected at an MOI of 0.5 with virus prepared using 90% decoy expression would be expected to integrate Perturb-seq constructs. Therefore, the decision to use (and how to use) a decoy-based approach will depend most strongly on the requirements and goals of individual experiments. Promisingly, an integration-defective decoy could be used to circumvent the problem of excess integration, as has been successfully piloted elsewhere (Feldman et al., 2018). However, hereto there are limitations; notably, libraries prepared with integration-defective decoys would remain fully infectious.

Another limitation of the decoy-based approach for Perturb-seq library preparation is that conditions for successful suppression of barcode uncoupling during transduction may need to be individually optimized for each library design and possibly every cell type evaluated. This is because lentiviral recombination rates can vary depending on features like the sequence and secondary structure of RNA genomes or dNTP availability (DeStefano et al., 1992; Negroni and Buc, 2001; Operario et al., 2006).

Promisingly, CROP-seq, an alternative platform for single-cell expression profiling of CRISPR-based screens, cleverly avoids the problem of barcode uncoupling altogether (Datlinger et al., 2017). This platform uses sgRNA protospacer sequences as unique identifiers instead of independent barcodes. To accomplish this, the CROP-seq transfer vector (CROPseq-Guide-Puro) encodes a single Pol III-controlled sgRNA expression cassette within the U3 region of the lentiviral 3ʹ LTR (Figure 2). This places the sgRNA protospacer close to the viral polyadenylation signal sequence and at the 3ʹ end of a Pol II-controlled, polyadenylated RNA that can be captured within scRNA-seq libraries. Encoding the sgRNA cassette in this position, however, from within a Pol II cassette, raises the possibility that promoter interference will repress expression of the active Pol III-driven molecule. This concern is consistent with low sgRNA activity that has been observed when using an sgRNA expression construct that incorporates overlapping Pol II and Pol III transcripts (Hill et al., 2018). However, CROPseq-Guide-Puro also avoids this concern. This is because the entire sgRNA-containing LTR is duplicated upstream of the Pol II promoter during provirus synthesis, thus producing a second copy of the expression cassette (* in Figure 1). Moreover, because LTR duplication is not subject to intermolecular recombination (discussed above), the design of CROPseq-Guide-Puro allows faithful transduction of co-packaged sgRNA libraries with matched protospacer “barcodes”. Recently, targeted PCR amplification of these “barcodes” was shown to improve guide identification in CROP-seq screens (Hill et al., 2018). As barcode amplification was first pioneered for Perturb-seq screens, this latest implementation represents a promising merger of the two platforms (Adamson et al., 2016; Dixit et al., 2016). Independently, we (and others) have begun using similar hybrid strategies to conduct experiments that warrant pooled library preparation.

As with all approaches discussed herein, the CROP-seq platform, with or without barcode amplification, has both advantages and limitations. As illustrated by Datlinger *et al*, positioning the sgRNA expression cassette within the lentivirus U3 region may place an upper limit on cassette size. This may restrict the use of CROPseq-Guide-Puro to delivery of single sgRNAs because combinatorial sgRNA delivery would require inserting multiple cassettes into a single LTR. Moreover, because recombination can disrupt sequences *within* LTRs (Yu et al., 1998), CROPseq-Guide-Puro presents no obvious advantage for library designs that incorporate tandem cassettes.

Moving forward, programmed delivery of combinatorial perturbations will be essential for many studies, especially those that seek to delineate and understand the role of genetic interactions. For this reason, Perturb-seq was designed from the beginning to enable the simultaneous transduction of up to three linked sgRNAs (and potentially more) without interference from intramolecular recombination (Adamson et al., 2016). Additionally, because Perturb-seq marks each set of linked sgRNAs with a single GBC, combinatorial guide assignment on this platform is straightforward. Although only tested for suppression of intermolecular scrambling among single, barcoded sgRNA expression vectors thus far, a decoy approach is likely also amenable for enabling faithful delivery of co-packaged combinatorial Perturb-seq vectors. However, for experiments of modest scale, we again consider arrayed lentiviral packaging a tractable option to limit intermolecular recombination.

An alternative strategy for delivering combinatorial perturbations to target cells is supertransduction of single sgRNA expression vectors. Here too, though, there are disadvantages. Perhaps most notably, pairing perturbations through random infection events will generate tails of cells that receive either one or greater than two sgRNAs, causing excess or unwanted sampling of single perturbations and higher-order interactions. Perturbation pairs could also be generated by simultaneous transduction of separately marked libraries that could be co-selected. However, limiting the copy number of each library while maximizing pairs would remain an experimental challenge. In either case, obtaining pairs at random may compound variation in sgRNA representation. Programmable combinations in contrast allow complete control. This means that uninteresting combinations can be avoided and those thought to have strong phenotypes can be systematically enriched.

In summary, as functional genomics studies have begun to require delivery of increasingly complicated transgene constructs, uncoupling of linked variable regions during lentiviral transduction has emerged as a key experimental challenge. Here, we have demonstrated two methods of limiting uncoupling in Perturb-seq screens, and we have reviewed considerations for designing successful single cell experiments. Moving forward, however, efforts to engineer more robust systems for transgene delivery could fill a general need. Notably, nature may lend insight to such efforts, as other retroviruses generate recombinants at lower frequencies than HIV-1. In the case of Moloney murine leukemia virus (MLV) this is likely due to preferential co-packaging of homotypic genomes (Flynn et al., 2004; Onafuwa et al., 2003; Zhuang et al., 2006). Alternatively, leveraging an RT less prone to template switching or forcing virions to packaging only single genomes could be explored. Nevertheless, in the end, the “best” design for experiments that leverage lentiviral transgene libraries will always balance the desire for efficiency and scale with individual research objectives.

## METHODS

### Plasmid design and construction

Transfer vectors pMJ002 and pBA613 were cloned from the Perturb-seq vector backbone (pBA439, Addgene #85967). For other purposes, 20 bp PCR primer-binding sites were added directly upstream and downstream of the modified mouse U6 (mU6)-controlled sgRNA expression cassette to make pMJ002. This vector, like pBA439, expresses an sgRNA programmed with the GFP-targeting EGFP-NT2 protospacer called sgGFP (Gilbert et al., 2013) and BFP. To create pBA613, the protospacer in pMJ002 was replaced with NegCtrl-3 (5ʹ- GACGACTAGTTAGGCGTGTA-3ʹ) (Adamson et al., 2016), and BFP was replaced with mCherry by Gibson assembly using synthesized DNA. The lentiDecoy vector (pBA889) was cloned from pInducer20 (Meerbrey et al., 2011). After digestion with PstI and AgeI to remove the attR1-CmR-ccDB-attR2-Ubc-rtTA3-IRES region, an HA-tag was inserted between the TRE2 promoter and Neo by Gibson assembly using synthesized DNA.

### Cell culture, viral transduction, and cell analysis

K562 cells expressing dCas9-BFP-KRAB and GFP under the control of the SV40 promoter (Adamson et al., 2016) were grown in RPMI-1640 with 25 mM HEPES, 2.0 g/L NaHCO3, 0.3 g/L L-Glutamine supplemented with 10% FBS, 2 mM L-Glutamine, 100 units/mL penicillin, and 100 µg/mL streptomycin. HEK293T cells were grown in Dulbecco’s modified eagle medium (DMEM) supplemented with 10% FBS, 100 units/mL penicillin, and 100 µg/mL streptomycin. HEK293T cells were transfected with transfer plasmids alongside standard packaging and envelope plasmids using *Trans*IT-LTI Transfection Reagent (Mirus, MIR2306) to produce lentivirus. Transfer plasmids pMJ002 and pBA613 were transfected individually or at roughly equimolar amounts. pBA889 (lentiDecoy) was co-transfected at 0, 56%, and 92% of total transfer plasmids. Two days later, virus was harvested, filtered through an 0.45 µM syringe filter, and precipitated with Lentivirus Precipitation Solution (ALSTEM, #VC100) according to manufacturer’s instructions. Virus was concentrated to 20x. K562 cells were infected with centrifugation at 1000 x *g* for 2 hours at 33°C. Infections using virus prepared without lentiDecoy were performed with 12.5 ul. Infections using virus prepared with 56% and 92% lentiDecoy were performed with 25 ul and 125 ul virus, respectively. Post-spinfection, cells were removed from viral supernatant. Cells were analyzed by flow cytometry after 8 days.

## ACKNOWLEDGEMENTS

We thank L.M. Sack and S.J. Elledge for helpful discussion. This work was funded by National Institutes of Health grants to J.S.W. (P50 GM102706, U01 CA168370, R01 DA036858) and M.J. (F32 GM116331). J.S.W. is a Howard Hughes Medical Institute Investigator. T.M.N. is a fellow (DRG-[2211-15]) and B.A. was an HHMI fellow of the Damon Runyon Cancer Research Foundation (DRG-[2182-14]).

